# Remote homolog detection places insect chemoreceptors in a cryptic protein superfamily spanning the tree of life

**DOI:** 10.1101/2023.08.30.555482

**Authors:** Nathaniel J. Himmel, David Moi, Richard Benton

**Affiliations:** Center for Integrative Genomics; Department of Computational Biology, Faculty of Biology and Medicine University of Lausanne, CH-1015, Lausanne Switzerland

## Abstract

Many proteins exist in the so-called “twilight zone” of sequence alignment, where low pairwise sequence identity makes it difficult to determine homology and phylogeny ^1, 2^. As protein tertiary structure is often more conserved ^3^, recent advances in *ab initio* protein folding have made structure-based identification of putative homologs feasible ^4–6^. However, structural screening and phylogenetics are in their infancy, particularly for twilight zone proteins. We present a pipeline for the identification and characterization of distant homologs, and apply it to 7-transmembrane domain ion channels (7TMICs), a protein group founded by insect Odorant and Gustatory receptors. Previous sequence and limited structure-based searches identified putatively-related proteins, mainly in other animals and plants ^7–10^. However, very few 7TMICs have been identified in non-animal, non-plant taxa. Moreover, these proteins’ remarkable sequence dissimilarity made it uncertain if disparate 7TMIC types (Gr/Or, Grl, GRL, DUF3537, PHTF and GrlHz) are homologous or convergent, leaving their evolutionary history unresolved. Our pipeline identified thousands of new 7TMICs in archaea, bacteria and unicellular eukaryotes. Using graph-based analyses and protein language models to extract family-wide signatures, we demonstrate that 7TMICs have structure and sequence similarity, supporting homology. Through sequence and structure-based phylogenetics, we classify eukaryotic 7TMICs into two families (Class-A and Class-B), which are the result of a gene duplication predating the split(s) leading to Amorphea (animals, fungi and allies) and Diaphoretickes (plants and allies). Our work reveals 7TMICs as a cryptic superfamily with origins close to the evolution of cellular life. More generally, this study serves as a methodological proof of principle for the identification of extremely distant protein homologs.

## Results and Discussion

Insect Odorant receptors (Ors) and Gustatory receptors (Grs) are 7-transmembrane domain ion channels (7TMICs) critical for the behavior and evolution of insects ^7,11,12^. Although originally thought to be insect-specific ^13–18^, the genomic revolution enabled sequence-based searches to identify putative homologs in animals (Gustatory receptor-like proteins; Grls), plants (DUF3537 proteins) and single-celled eukaryotes (GRLs) ^7–9,19^. However, the representation of 7TMICs across taxa remained sparse, recognized in only a small number of unicellular eukaryotes (17 proteins from 7 species), and missing from several holozoan lineages, including chordates, choanoflagellates, comb jellies and sponges ^7–9,19^.

The best-characterized 7TMICs are insect Ors, which function as odor-gated heterotetrameric (or in some cases homotetrameric) ion channels ^20–23^. A substantial breakthrough came from two Or cryo-electron microscopy structures: the fig wasp *Apocrypta bakeri* Or co-receptor Orco ^20^ and the jumping bristletail *Machilis hrabei* Or5 ^21^ (**Figure 1A**). Or monomers have several notable structural features, including: (i) 7 transmembrane alpha helices with a characteristic packing pattern; (ii) an intracellular N-terminus and extracellular C-terminus; (iii) shorter extracellular than intracellular loops; (iv) long TM4, TM5, and TM6 helices that extend into the intracellular space, forming the “anchor domain,” where most inter-subunit interactions occur; (v) an unusual “split” TM7 helix, composed of an intracellular TM7a (part of the anchor domain) and a transmembrane-spanning TM7b (which lines the pore of the ion channel); and (vi) an N-terminal re-entrant loop (TM0) ^20,21,24,25^. These tertiary structural features are remarkably highly-conserved despite low primary sequence conservation; for example, the two experimental structures have virtually indistinguishable folds while having only 19% amino acid sequence identity (**Figure 1B**). Importantly, these structures can be accurately predicted *in silico* by several algorithms ^8, 25^, notably AlphaFold (**Figure 1C**) ^4, 10^.

**Figure 1.**
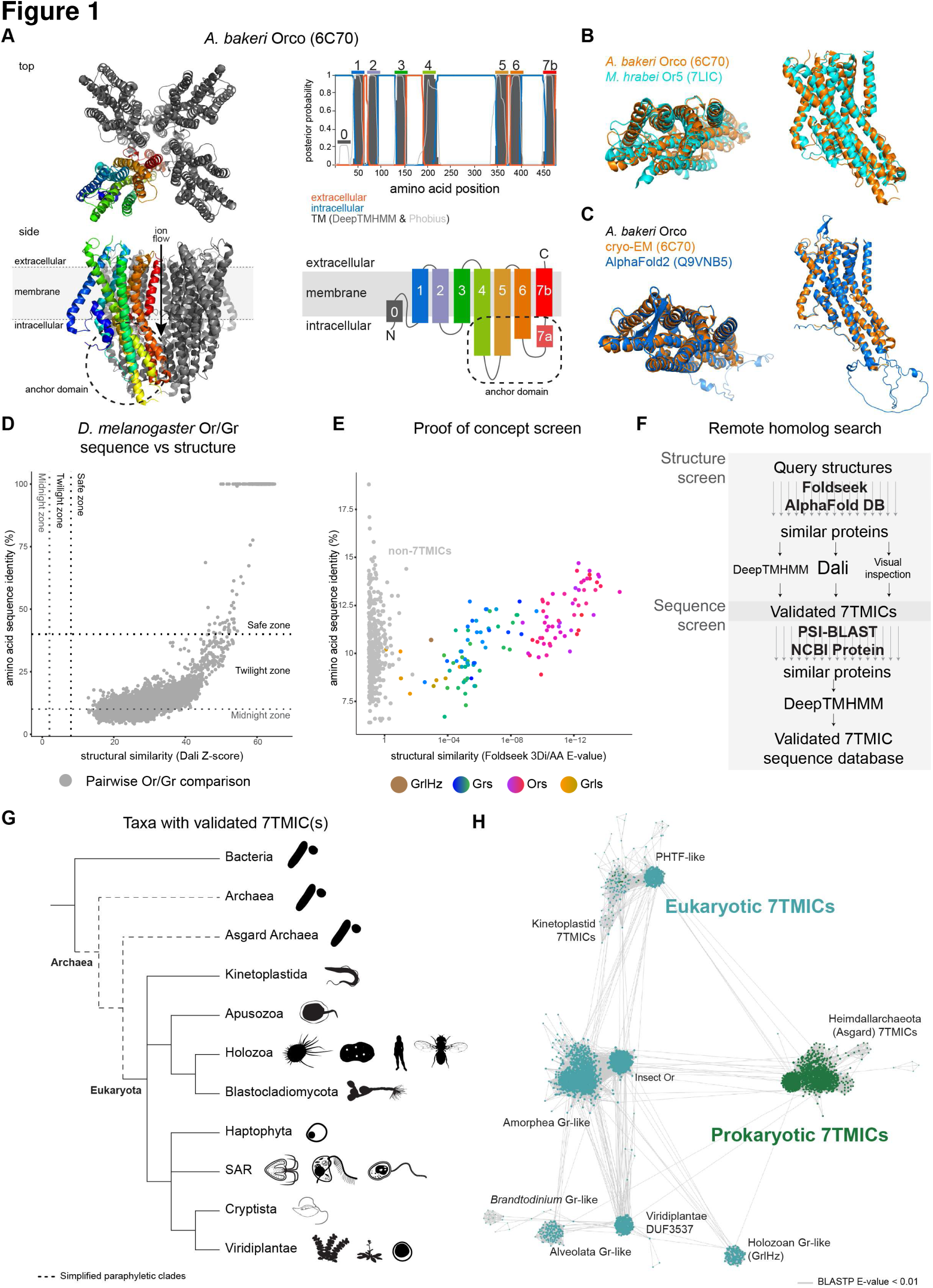
A structure– and sequence-based screen for the identification and validation of extremely distant 7TMIC homologs. (**A**) Left: top and side views of the cryo-EM structure of the *A. bakeri* Orco homotetramer (PDB 6C70), with one subunit colored ^20^. Right: transmembrane prediction of *A. bakeri* Orco by DeepTMHMM and Phobius illustrating the characteristic membrane topology of 7TMICs. A cartoon representation of 7TMIC membrane topology is shown below. (**B**) Aligned cryo-EM structures of *A. bakeri* Orco and *M. hrabei* Or5 (PDB 7LIC). (**C**) Aligned cryo-EM and AlphaFold structures of *A. bakeri* Orco. (**D**) Sequence identity versus structural similarity for all pairwise comparisons of AlphaFold models of *D. melanogaste*r Ors and Grs, using Dali. The cluster of dots at the top right are self-to-self comparisons and isoforms of the same gene. (**E**) Proof of principle Foldseek screen of the AlphaFold structural proteome of *D. melanogaster*, with results of the screen plotted by Foldseek-derived percent amino acid sequence identity and E-values. (**F**) Outline of the screen and validation pipeline. (**G**) Cladogram of taxa in which 7TMICs were identified. (**H**) All-to-all BLASTP network of 7TMICs (each represented by a dot), which visualizes only pairwise sequence similarity. Several clusters form, suggesting monophyly within clusters (annotated manually based on CLANS clustering). At presumed longer evolutionary distances, 7TMICs show little-to-no pairwise sequence identity, represented by the weak connectivity (i.e. few edges) between most clusters.

Recently, we took advantage of the structural similarity of 7TMICs to perform structure-based screens for putative homologs that had not been identified by sequence-based screening. These screens identified several proteins adopting the 7TMIC fold, including: fly-specific Gustatory receptor-like proteins (Grls); a highly-conserved lineage of eukaryotic proteins (PHTFs, an acronym for the misnomer Putative Homeodomain Transcription Factor); a holozoan-specific Grl lineage (GrlHz); and trypanosome 7TMICs ^10^. However, these searches were limited by the high computational requirements of the structural alignment tool—Dali ^26, 27^—and only ∼564,000 AlphaFold models from 48 species were screened. Thus, large taxonomic gaps still exist: fewer than 50 proteins have been identified outside of animals and plants, and none have been identified in prokaryotes (despite screening 17 prokaryotic proteomes ^10^). Beyond the technical limitations leading to sparse taxonomic sampling, the PHTF, GrlHz, Gr/Or, DUF3537 and various unicellular eukaryotic 7TMIC proteins share little to no recognizable sequence similarity. It is thus unclear how many 7TMICs exist across taxa and if 7TMICs form a single or many homologous protein families. We thus sought to build a new pipeline for remote homolog detection, validation, and sequence/structure analysis, aiming to resolve the evolutionary history of 7TMICs, be they homologous or convergent.

### Insect Ors and Grs have high structural similarity despite exceptional sequence dissimilarity

Comparisons of the AlphaFold models of *Drosophila melanogaster* Ors and Grs exemplifies the discordance between sequence and structure similarity: pairwise comparisons average only ∼13% pairwise amino acid sequence identity (**Figure 1D**, y-axis), placing these proteins at the border of the so-called “twilight zone” (10-40% sequence identity) ^1^ and “midnight zone” (<10% sequence identity) ^2^ of sequence alignment. By contrast, pairwise comparisons of the corresponding AlphaFold structures—using Dali Z-scores, a widely-used metric of fold similarity ^26, 27^—reveals that all pairwise comparisons fall within the “safe zone” of structural alignments, indicating high statistical confidence in their similarity (**Figure 1D**, x-axis). When visualized as a sequence similarity network (produced by all-to-all BLASTP searches), Ors and Grs—together with other *D. melanogaster* 7TMICs, i.e. Grls and Phtf ^10^—segregate into several non-contiguous clusters (**Figure S1A**). This analysis demonstrates that no single receptor protein can be used to identify all others via simple sequence-based searches. By contrast, structure-based search strategies (e.g. Dali, **Figure S1B**) are capable of densely networking these proteins. As *D. melanogaster* Ors and Grs are just a very small subset of 7TMICs that likely had a single common ancestor ^28^, these observations emphasise how structure-based screens are a greatly superior way to search for distant homologs across more phylogenetically diverse species ^3^.

### A pipeline for identifying extremely distant protein homologs

Foldseek—a recently released tool for structure-based protein comparisons— operates orders of magnitude faster than Dali and other structural alignment tools, making large protein homolog screens feasible ^5^. We first benchmarked Foldseek on *D. melanogaster* 7TMICs. When forced to compare the AlphaFold model of Orco to all other 7TMICs of this species, Foldseek produced structural similarity scores that correlate with Dali Z-scores (**Figure S1C**). As proof of concept for screening, we used the *D. melanogaster* Orco AlphaFold model to survey the AlphaFold structural proteome of *D. melanogaster* (**Figure 1E**). Foldseek was able to recover all *D. melanogaster* 7TMICs except Phtf: thus, the method can result in false negatives. However, with the most permissive settings—which would allow the most sensitive homolog detection—Foldseek also had an extremely high false positive rate (73.3%), and the most divergent relatives (e.g. Grls) had higher E-values and/or lower percent sequence identity than false positives (**Figure 1E**). As we were interested in screening for distant and divergent 7TMIC homologs across much longer evolutionary distances than only within *D. melanogaster*, we recognized that neither E-value nor sequence identity could serve as an effective threshold. These benchmarks illustrated the need for additional search and validation steps to minimise both false positive and false negative results in our screen.

We therefore implemented Foldseek as part of a screening and validation pipeline, with the goal of determining the presence or absence of 7TMICs across the tree of life (**Figure 1F**). This pipeline first uses Foldseek to search for structurally similar models in the AlphaFold Protein Structure Database, which currently consists of ∼200,000,000 models from >1,200,000 species (see Methods for details on exclusions). After structural validation, it employs PSI-BLAST in a sequence-based screen, providing structurally-informed access to >400,000,000 sequences—with diverse transcriptomic, proteomic, genomic, and metagenomic origins—that might not have a corresponding protein model. This second step also allows for the identification of proteins with models that were missed in the first structure-based screen, which we expected to occur due to the occurrence of false negatives at hypothetically vast evolutionary distances (e.g. Orco to Phtf (**Figure 1E**)).

As false positives can have high scores, and as some public data can be incomplete or of low quality, we implemented several verification steps to extract true hits. For proteins identified by structural model, we: (i) curated proteins based on membrane topology as predicted by the protein language model DeepTMHMM ^29^; (ii) validated structural alignments using Dali; and (iii) visually inspected putative hits for the previously-described 7TMIC features. Proteins identified through sequence similarity were curated based on membrane topology (DeepTMHMM).

### 7TMICs are present across the tree of life

This screen recovered thousands of previously unidentified 7TMICs spanning the tree of life (**Figure 1G-H** and **Figure S1E**). These hits not only include new eukaryotic 7TMICs (hereafter, Euk7TMICs), but also sequences from all major branches of bacteria (Bac7TMICs) and archaea (Arch7TMICs) (see “Protein nomenclature” section in the Methods). These proteins come from several obviously monophyletic clades, apparent as clusters in a network representing all-to-all BLASTP searches (**Figure 1H**). However, they can exhibit very little pairwise sequence similarity, represented by few edges between clusters in the BLASTP network (**Figure 1H**).

Euk7TMICs could be visually sorted into two types of structure: Or/Gr/Grl/GRL/DUF3537-like (**Figure 2A**), having the canonical insect Or-like fold; or PHTF-like (**Figure 2B**), having the same core structure, but with a long first intracellular loop (IL1). While the various prokaryotic 7TMICs have a striking degree of structural similarity to Euk7TMICs (**Figure 2C-E**), we observed that they generally had shorter TM4 and TM5 helices, which constitute a component of the anchor domain in insect Ors (**Figure 1A**). Heimdallarchaeota 7TMICs (**Figure 2E**) were an exception: their overall tertiary structure appeared eukaryote-like. This qualitative similarity (supported by subsequent quantitative analyses, described below) is notable, as Heimdallarchaeota are proposed to be the most closely related extant archaea to eukaryotes ^30–34^. In addition, a small number of metagenomically-identified prokaryotic 7TMICs have Euk7TMIC-like folds (**Figure 2F**). Notably, these show high sequence similarity to Euk7TMICs (green nodes in the eukaryotic PHTF-like cluster, **Figure 1H**), suggesting that these sequences are the result of eukaryote-to-prokaryote horizontal gene transfer(s), a hypothesis further supported phylogenetically (see below).

**Figure 2.**
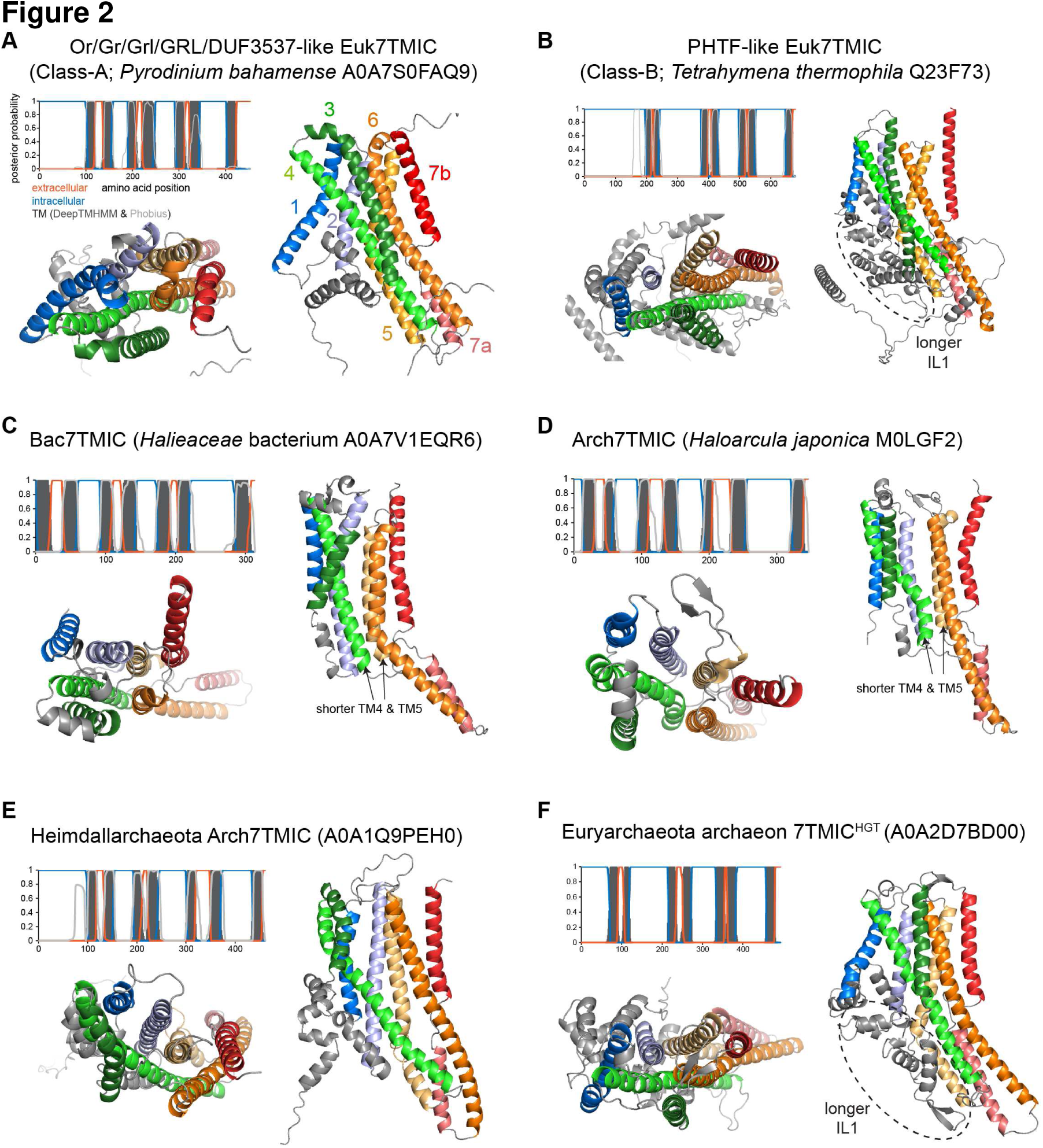
Examples of newly-identified 7TMICs. Transmembrane predictions, and top and side views of the AlphaFold structure of newly-identified 7TMICs. (**A**) Representative example of a Grand DUF3537-like Euk7TMIC, subsequently phylogenetically classified as Class-A (Figure 4). These proteins have all the stereotyped 7TMIC features. (**B**) Representative example of a PHTF-like Class-B 7TMIC (Figure 4). These proteins have stereotyped 7TMIC features, with the addition of a long intracellular loop between TM2 and TM3 (IL1). (**C-E**) Representative examples of a Bac7TMIC and Arch7TMICs. When clustered by Foldseek at 90% coverage (data not shown), prokaryotic 7TMICs form three structure clusters (represented here in (C), (D), and (E)), although they cannot be easily distinguished visually. These proteins all share the stereotyped 7TMIC features but, with the exception of Heimdallarchaeota 7TMICs (E), have shorter TM4 and TM5. (**F**) Representative example of the small number of bacterial and archaeal proteins with extreme fold and high sequence-similarity to Euk7TMICs, which are presumed to have arisen through horizontal gene transfer(s) (HGT) (Figure 4).

### 7TMICs have a shared tertiary structure and amino acid sequence profile, supporting homology

While we observed structural similarities between the proteins our screen identified, it remained unclear if these sequences are homologous, or if they represent cases of structural convergence. To address this fundamental issue, we adapted established protein comparison tools into a graph-based approach for determining homology based on both structure and sequence. For protein structures, we calculated all-to-all template modelling (TM) scores, where those >0.5 indicate high statistical confidence of fold similarity ^35^. For protein sequences, we performed all-to-all PSI-BLAST searches; PSI-BLAST builds iterative multiple sequence alignments, thereby identifying distant homologs by family-wise sequence profiles, rather than by simple pairwise sequence similarities ^36^. In essence, PSI-BLAST networking is equivalent to performing PSI-BLAST homolog searches starting with every structurally-validated 7TMIC as a query (see Methods). For both methods, one expects homologous proteins to form bidirectional connections between each other (i.e. that pairs will be reciprocal hits), and that homologous families will be highly interconnected, thereby collapsing into visually identifiable clusters in structure– and sequence-space. We performed these analyses with Type-I and Type-II opsins as control groups, as these large families are 7-transmembrane domain proteins (unrelated to 7TMICs) that adopt highly similar folds to one another, despite no recognized sequence similarity ^37^.

In the structural similarity network, 7TMICs formed a densely connected linkage cluster, disconnected from a unified opsin linkage cluster (**Figure 3A**). 7TMICs also clustered in sequence space – after 3 PSI-BLAST iterations, 7TMICs collapsed into a single, highly connected community structure (**Figure 3B** and **Figure S2A**). In stark contrast, the opsins separated into distinct Type-I and Type-II community structures, demonstrating that structure and sequence are not necessarily linked (**Figure 3B**). While there were connections between 7TMICs and the opsins, in the third iteration these constituted only 18 of the 1,117,609 connections (0.0016%), almost certainly representing spurious similarity. A small minority of 7TMICs (33/2421 representative sequences) from diverse eukaryotic taxa showed no connectivity to the core 7TMIC cluster in the second iteration and weak connectivity in the third; these may be extremely rapidly evolving proteins and/or cases of independent structural convergence.

**Figure 3.**
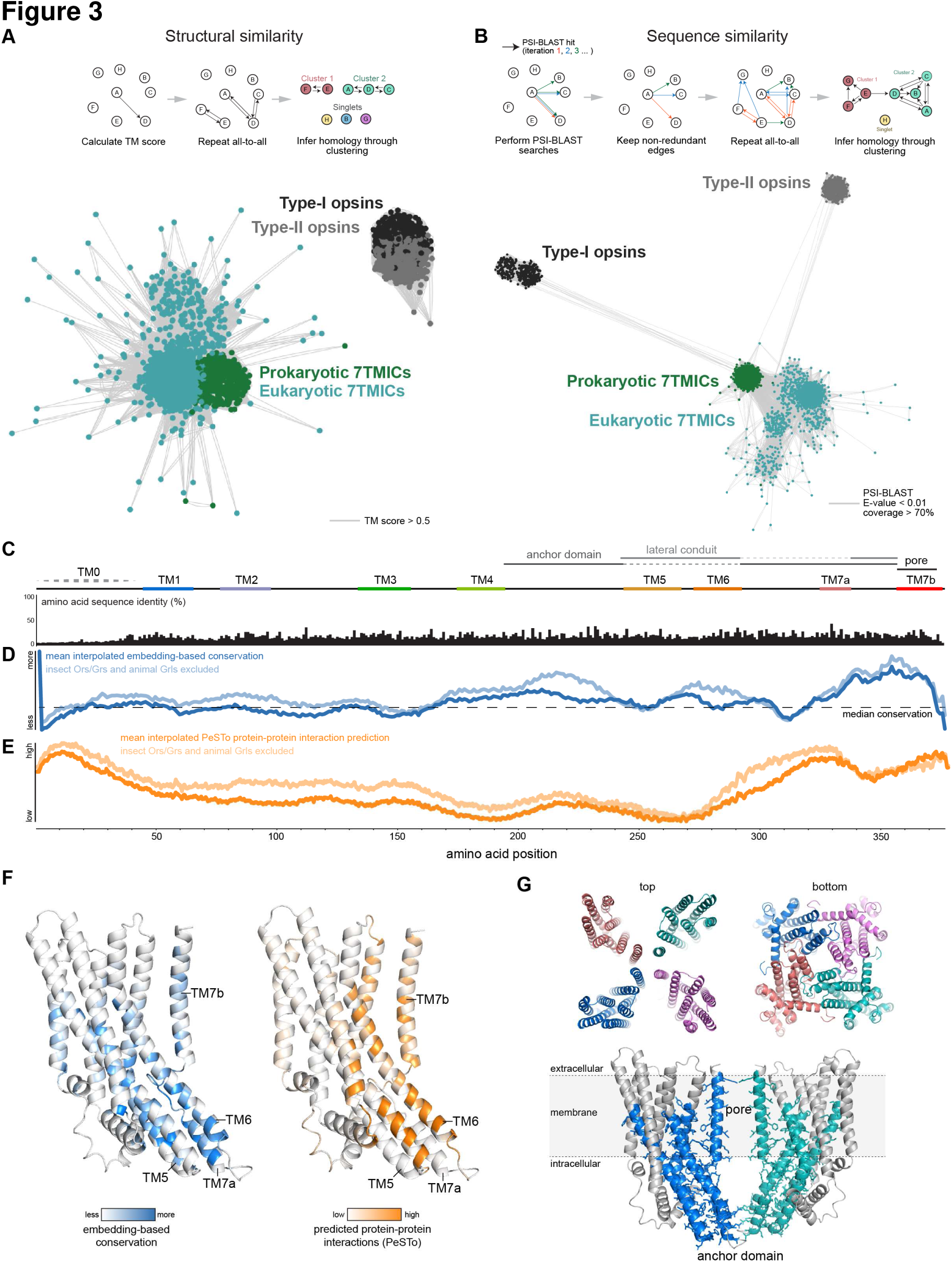
Evidence for 7TMIC homology through structural and sequence similarity. (**A**) Structural similarity network of 7TMICs derived from all-to-all TM-scores (schematized at the top). (**B**) Sequence similarity network of 7TMICs produced by all-to-all PSI-BLAST searches (schematized at the top; iteration 3 is shown), providing evidence that 7TMICs have a family-wide sequence profile. This pattern of strong, bidirectional linkages became apparent already in PSI-BLAST iteration 2, while subsequent iterations resembled iteration 3 (**Figure S2A** and Supplemental Material). (**C**) Amino acid sequence identity derived from a query-centered Foldseek alignment for the centermost node in the structural similarity network (*Symbiodium natans* A0A812K102). Transmembrane predictions are from Phobius. TM7b was annotated manually. TM0 (the re-entrant loop) is indicated with a dashed line, as it is inconsistently predicted by DeepTMHMM and Phobius, and often predicted with low confidence region in AlphaFold models. Query-centered alignments for all 7TMIC models analyzed here are available in the supplemental data. (**D**) Average sequence embedding-based conservation scores for 7TMICs, with the curves interpolated to match the length of A0A812K102. Column conservation scores are significantly correlated with column sequence identity for A0A812K102 (**Figure S2B**). The location and strength of conservation likely varies by 7TMIC family and subfamily, and excluding the well-established 7TMICs (insect Ors/Grs and animal Grls) led to overall increased conservation scores (light blue line, also **Figure S2C**). Embedding-based conservation score for all 7TMIC models analyzed here are available in the supplemental data. (**E**) Average PeSTo protein-protein interaction predictions. The region with the most consistently-predicted protein-protein interactions is near the C-terminus, correlating with the site of highest sequence conservation; again, exclusion of Ors/Grs/Grls led to higher prediction scores (light orange line). PeSTo predictions for all 7TMIC models analyzed here are available in the supplemental data. (**F**) Sequence-embedding based conservation scores (blue) and PeSTo-derived protein-protein interaction scores (orange) mapped onto the AlphaFold model of A0A812K102. (**G**) Top: top (presumed extracellular) and bottom (presumed intracellular) views of a hypothetical tetramer of A0A812K102 (predicted by AlphaFold-Multimer), showing that individual subunits have their closest interactions in the pore and anchor domains, similar to Ors ^20, 21^. Bottom: side view of the A0A812K102 tetramer, with two subunits masked for clarity, and the presumed anchor and pore regions colored on the visualized subunits. In total, we modelled 85 tetramers; 83 of these were Or-like, in that they displayed rotational symmetry, with the closest interactions in the hypothetical anchor and pore regions (further examples in **Figure S2D**).

We next sought to determine which, if any, regions of 7TMICs are more conserved. It was previously observed that insect Ors display the highest conservation in the anchor domain and pore-forming region, with greater divergence in the N-terminal region that forms the odor-binding pocket ^20, 21^. We calculated sequence embedding-based conservation scores, which identify sites that are evolutionarily constrained ^38^. This analysis elucidated a similar conservation pattern for newly identified 7TMICs: while absolute amino acid sequence identity is low (averaging 15% across sites, **Figure 3C** and **Figure S2B**), embedding-based conservation analysis revealed that the most highly conserved regions are in three locations: the hypothetical anchor domain (intracellular sequences spanning TM4-TM5 and TM6-TM7a), the hypothetical pore (TM7b), and TM5-TM6, which form lateral ion permeation conduits in Ors ^20, 21^ (**Figure 3D**, **3F**). We next used the protein language model PeSTo ^39^ to predict protein-protein interactions in 7TMICs, revealing two conserved regions (**Figure 3E, 3F**). The first was N-terminal, corresponding to the re-entrant loop (TM0); this region has an important, albeit poorly-understood, function in Orco ^25^. The second region was in the hypothetical anchor domain and pore, in the same regions as the highest peaks of sequence conservation.

These findings are not biased by the inclusion of proteins previously determined to be homologous (insect Ors/Grs and animal Grls); on the contrary, removing these sequences improved average conservation (and interaction) scores in these regions (**Figure 3D-E** and **Figure S2C**).

While we cannot know *a priori* whether these proteins form tetramers like insect Ors, these patterns of conservation and predicted protein-protein interactions suggests they may assemble as multimers using the same domains. Consistent with this idea, using AlphaFold-multimer ^40–43^ to predict complexes of tetramers of newly-identified 7TMICs, the vast majority of resulting quaternary structures had striking similarity to experimentally-derived Or structures (**Figure 3G** and **Figure S2D**). In these models, the hypothetical anchor domain (particularly TM7a) contains the closest protein-protein interactions and TM7b lines the putative pore.

These results quantitatively demonstrate that 7TMICs have a common structure, a shared sequence profile, and similar patterns of sequence conservation. Thus, the most parsimonious hypothesis is that 7TMICs are a homologous protein superfamily.

### The evolutionary history of 7TMICs

Having obtained evidence for the homology of 7TMICs, we next sought to elucidate the evolutionary history of the superfamily. As we expected pairwise sequence dissimilarity would make multiple sequence alignments difficult, we performed sequence-based phylogenetics on an ensemble of alignments, thus resulting in a “forest” of phylogenetic trees (**Figure S3A-E**), from which we extracted the median sample tree (**Figure 4A**). These analyses suggested that there are two main Euk7TMIC families, hereafter termed Class-A and Class-B Euk7TMICs. While Class-A Euk7TMICs appear to be monophyletic, the monophyly of Class-B is uncertain. Class-A Euk7TMICs include insect Ors/Grs, animal Grls, plant DUF3537, holozoan GrlHz, and various unicellular eukaryotic 7TMICs. Class-B Euk7TMICs are PHTF-like proteins from diverse taxa, including a small number of bacterial and archaeal proteins (see green nodes in the PHTF-like cluster the BLASTP network (**Figure 1H**) and an example structure (**Figure 2F**)). The phylogenetic separation of these from other prokaryotic 7TMICs suggests they arose through horizontal gene transfer(s). This analysis also suggests that kinetoplastid 7TMICs (Kineto7TMICs) branch more proximally to prokaryotic 7TMICs, consistent with the hypothesis that kinetoplastids (and allies; collectively Discoba) split early in eukaryotic evolution ^44^. The median sampled tree (**Figure 4A**) generally represents this diverse tree space: Kineto7TMICs branch proximally to Arch/Bac7TMICs (here deeply, but with low branch support; 0.79 and 0.409 for the two most proximal branches); Class-A is monophyletic, with modestly strong branch support (0.91); and Class-B is paraphyletic, but with extremely low branch support on the relevant branch (0.22) (**Figure 4A**).

**Figure 4.**
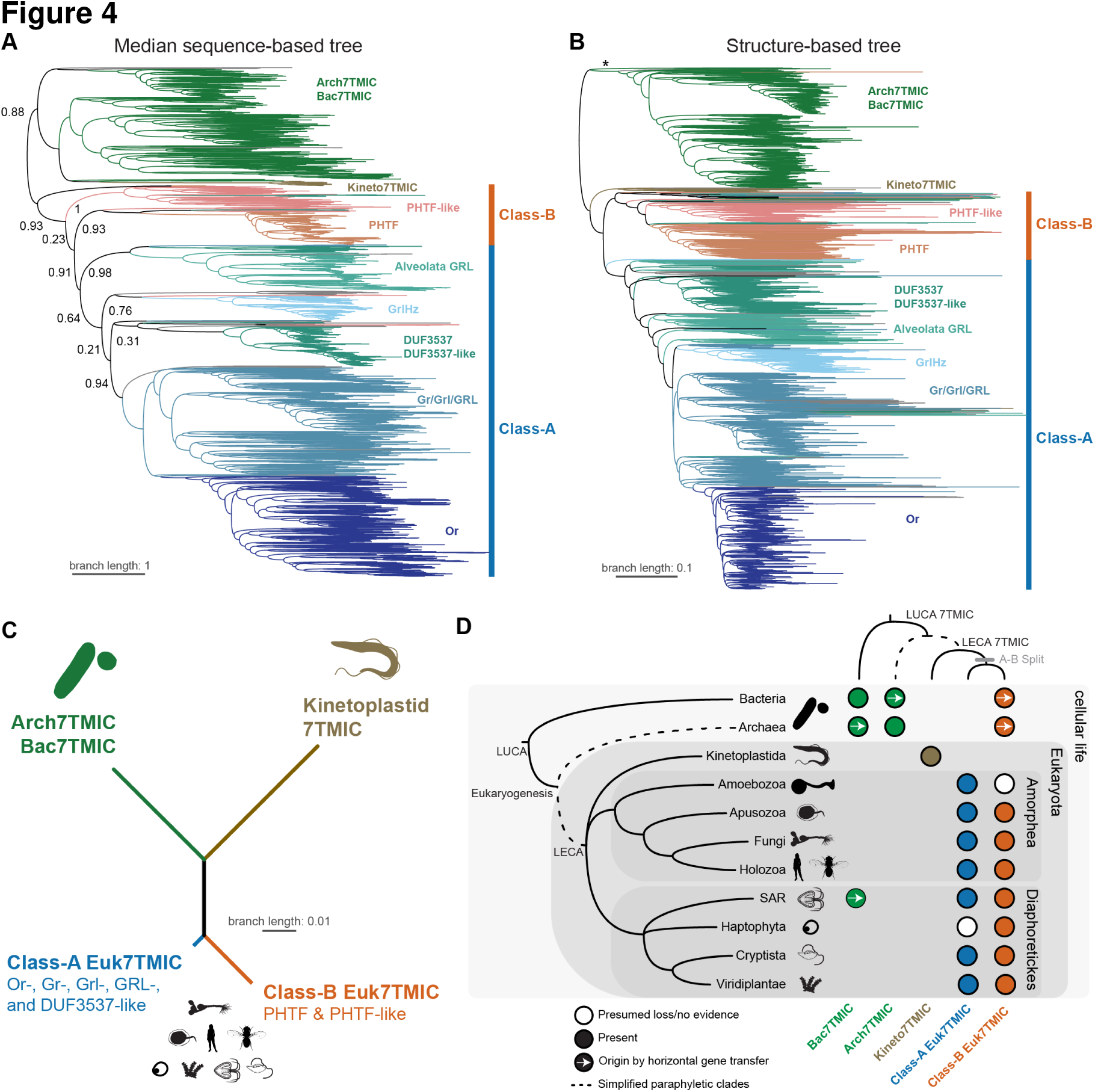
A model for the evolution of the 7TMIC superfamily. (**A**) The median phylogenetic tree of 7TMICs sampled from the Robinson-Foulds-based tree space of 48 sequence-based phylogenetic trees. For visualization purposes, the tree is arbitrarily rooted in the last common ancestor of all Arch/Bac7TMICs (which are highly reticulated); the true root is likely at the unidentified location of the last universal common ancestor (LUCA) within the prokaryotic branch. Branch lengths are derived from the average number of substitutions per site. Tree space is visualized in **Figure S2C-E**, and all trees and alignments are available in the Supplemental Data. (**B**) TM-score based structural tree of 7TMICs derived from fold_tree, automatically rooted using the MAD method. As in (A), the true root is likely at the unidentified location of LUCA. Branch lengths are derived from the underlying distance matrix of TM-scores. The asterisk marks the branch containing Heimdallarchaeota 7TMICs; a fully annotated tree is available in the supplemental data. (**C**) Collapsed version of the tree in (B) highlighting the major branching patterns. Kinetoplastid 7TMICs likely branched early, while the Class-A/Class-B split occurred after the emergence of the last eukaryotic common ancestor (LECA), but before the split(s) leading to Amorphea and Diaphoretickes. (**D**) Summary of the results of the screen and evolutionary analyses. The left tree shows assumed relationships between the various taxa in which 7TMICs were identified, while the top tree shows the evolutionary history of 7TMICs themselves. The colored dots represent the presence or absence of 7TMIC families. At the subfamily level, many Class-B PHTF-like proteins may be the result of horizontal gene transfer(s), as there is broad but sparse taxonomic diversity within this putative subfamily. Note that this screen did not recover previously identified 7TMICs from Amoebozoa or Chytridiomycota, which were inferred to be Class-A based on previously described sequence similarity to insect Grs/Ors ^8^; in addition, here, “Fungi” only refers to Chytridiomycota and Blastocladiomycota.

To complement the sequence-based phylogenetic approach, we also employed a recently-developed structure-based phylogenetic method (fold_tree), which infers a minimum evolution tree from a matrix of Foldseek-derived structural alignments ^45^. We made three notable observations of the resulting tree (**Figure 4B**), which shared many similarities to the sequence-based phylogenies (**Figure 4A**). First, the prokaryotic branch most proximal to the Euk7TMICs included Heimdallarchaeota 7TMICs (**Figure 4B**, asterisk), consistent with their proposed relation to eukaryotes, and suggesting that the shared Euk7TMIC structure (i.e. longer TM4 and TM5 (**Figure 2**)) emerged just before eukaryogenesis. Second, Kineto7TMICs were placed as the sister clade to all other eukaryotic 7TMICs, consistent with their presumed early branching ^44^. Third, Class-B Euk7TMICs were essentially monophyletic.

The most parsimonious interpretation of these data is that the Class-A/Class-B split is the result of a gene duplication which occurred after eukaryogenesis, but before the speciation event(s) leading to Amorphea and Diaphoretickes (**Figure 4C-D**).

### Concluding remarks

We have described a structure– and sequence-based screening strategy for identifying extremely distant transmembrane protein homologs, revealing that 7TMICs are present across the tree of life, including novel discoveries of representatives in Bacteria and Archaea. We have also shown that, despite substantial pairwise sequence dissimilarity, 7TMICs have extremely high structural similarity and identifiable family-wise sequence similarity. Together our results provide the first strong evidence that these disparate proteins form a single, homologous superfamily. This finding contrasts with the Type-I and Type-II opsins, whose structural similarity to each other might represent a case of convergent evolution ^37, 46^. Despite the phylogenetic breadth and conserved structure of 7TMICs, our knowledge of their function is almost entirely restricted to a subset of insect proteins ^47, 48^, which represents only a single, insect-specific lineage of this family. Our work lays a foundation for the analysis of the presumably diverse functions of 7TMICs across a wide range of species. Moreover, we suspect this ancient and cryptic superfamily is only one of many that wait to be discovered in the depths of the twilight zone of sequence space.

## Supporting information

Key resource table

## Acknowledgements

We thank Matteo Dal Peraro, Lucien Krapp and members of the Benton laboratory for comments on the manuscript, and Christophe Dessimoz for support of this work. Silhouettes in Figures 1 and 4 were sourced from PhyloPic (www.phylopic.org/). N.J.H. is supported by a Human Frontier Science Program Long-Term Postdoctoral Fellowship (LT-0003/2022L). D.M. is supported by a Swiss National Science Foundation grant (216623) to Christophe Dessimoz. Research in R.B.’s laboratory is supported by the University of Lausanne, an ERC Advanced Grant (833548) and the Swiss National Science Foundation (310030B-185377).

## Author Contributions

Conceptualization, NJH and RB; Methodology, NJH; Software, NJH and DM; Validation, NJH and DM; Formal analysis, NJH and DM; Investigation, NJH; Resources, NJH; Data Curation, NJH and DM; Writing – Original Draft, NJH; Writing – Review & Editing, NJH, DM, and RB; Supervision, RB; Project administration, NJH and RB; Funding acquisition, NJH and RB.

DM designed and performed the fold_tree analysis. NJH performed all other formal analyses.

## STAR Methods

### Resource availability

#### Lead contact

Further information and requests for resources and reagents should be directed to and will be fulfilled by the lead contacts, Nathaniel Himmel and Richard Benton.

### Materials availability

This study did not generate new unique reagents.

### Data availability

All data have been deposited in Dryad (https://doi.org/10.5061/dryad.fqz612jz9) and are publicly available as of the date of the publication.

## Method details

### Structural screen and validation

Proof-of-concept screens were carried out using local implementations of Foldseek ^5^ and DaliLite ^26, 27^. Subsequent structure-based screens of the AlphaFold Protein Structure Database (https://alphafold.ebi.ac.uk/) ^4, 6^ were performed on the Foldseek server (https://search.foldseek.com/), using the following query structures/models: *A. bakeri* Orco (6C70); *M. hrabei* Or5 (7LIC); *D. melanogaster* GrlHz (Q9W1W8); *B. belcheri* GrlHz (A0A6P5ACQ6); *T. adhaerens* GrlHz (B3RTY0); *Z. mays* DUF3537 (A0A1D6LEW8, B4FJ88, and B6SUZ0); *P. patens* DUF3537 (A0A2K1ICX7, A0A2K1JKU0, and A0A2K1L324); *D. melanogaster* Phtf (Q9V9A8); *H. sapiens* PHTF1 (Q9UMS5) and PHTF2 (Q8N3S3); *P. halstedii* PHTF (A0A0P1B782); *L. infantum* GRL1 (A4HWQ9); and *T. brucei brucei* GRL1 (Q57U78) (**Figure S1D**). For the initial screen, we masked the WD40 repeats in trypanosome GRL1 and the long intracellular loop 1 in PHTF, thus restricting the search to the core 7TMIC domain. We did not set a statistical threshold (E-value) for putative homolog identification. For eukaryotic hits, we initially considered all hits from the screen. For archaeal and bacterial hits, we took the more stringent approach of only further analyzing those that were hits for all the query groups (annotated in **Figure S3A**). We did not formally screen animal or vascular land plant species because we considered that these taxa have been sufficiently screened ^7–10,19^, and we were most interested in the very early evolution of 7TMICs. Indeed, preliminary Foldseek screens did not elucidate any obvious new plant– and/or animal-specific 7TMICs (data not shown).

Subsequent validation was performed in several steps. First, transmembrane topology was predicted using DeepTMHMM (the BioLib implementation at https://dtu.biolib.com/DeepTMHMM/ and a local implementation) ^29^. For putative eukaryotic homologs, we assessed these predictions alongside structural models (visualized in PyMol), looking for: (i) 7 predicted transmembrane alpha helices; (ii) shorter extracellular than intracellular loops; (iii) an intracellular N-terminus and extracellular C-terminus; (iv) longer TM4, TM5, and TM6 helices; and (v) the exceptional “split” TM7 helix ^20,21,24,25^. We did not consider the re-entrant loop (TM0) as a criterion, as it is inconsistently predicted by transmembrane prediction methods ^8, 10^. For archaea and bacteria, we only further assessed hits with exactly 7 predicted transmembrane segments in the stereotyped architecture. We also used Phobius (https://phobius.sbc.su.se/) ^49, 50^ and the transformer model PeSTo (https://pesto.epfl.ch/) ^39^ to predict transmembrane topology; both were used for visualization, but neither was used to curate sequences. Finally, we used a local implementation of DaliLite to compare all remaining hits with the original query structures and three negative controls. For the negative controls we selected an Adiponectin receptor (*Homo sapiens* ADPR1; 5LXG) and a channelrhodopsin (*Chlamydomonas reinhardtii* Channelrhodopsin-2; 6EID) in advance of the screen, as both have 7 transmembrane domains but are unrelated to 7TMICs; we added the ABC transporter permease (*Escherichia coli* A0A061Y968) *post hoc,* as many of the screen hits were errantly annotated as ABC transporters. Only hits with Dali Z-scores >8 as compared to 7TMIC queries were further analyzed. This threshold is based on Holm’s criteria ^51^, where Z-scores >20 indicate definite homology, 8-20 probable homology, 2-8 a “gray area” (here, “twilight zone”) and <2 non-significant (here, “midnight zone”). We conceptualized these scores as “protein fold similarity” in place of “homology,” as we infer homology based on a holistic view of sequence, structure, and taxonomic features. Pearson’s correlation analysis and the Bayesian equivalent were performed in JASP (https://jasp-stats.org/).

### Sequence-based homolog identification

For putative eukaryotic homologs, the results of the Foldseek screen were used to select query sequences. CLANS was used to generate an all-to-all BLASTP network (E-value cutoff 0.01), which was subsequently clustered by the global network clustering option ^52–54^. PSI-BLAST homolog searches were carried out using all singlets and a representative sequence from each cluster (the node with the highest neighborhood connectivity). Searches were run on the NCBI server (https://blast.ncbi.nlm.nih.gov/Blast.cgi) against the clustered non-redundant (clustered_nr) sequence database, until convergence. PSI-BLAST searches were performed with an E-value cutoff of 0.05, but final candidates were selected only if they had a minimum coverage of 50% (with coverage of the transmembrane region) and a final E-value at or below 10^-10^. For searches recovering canonical animal Grs/Ors/Grls, the PSI-BLAST searches were stopped when the top 1000 hits were recovered, as these searches quickly converged on tens of thousands of predominately insect sequences, which was computationally time-consuming and methodologically unnecessary for this study.

For Arch7TMIC homologs, sequence databases were likewise assembled using PSI-BLAST, using each of the structural screen hits as a query sequence. Compared to the eukaryote-based searches, we took a more stringent approach, setting an E-value cutoff of 10^-10^ for both the PSI-BLAST search and final hit selection. Query sequences that were orphans, or which had very few sequence-based homologs (<10), were excluded from further analyses. These searches recovered the Bac7TMICs, so efforts were not repeated using Bac7TMIC queries.

After preliminary homolog identification, DeepTMHMM was used to predict transmembrane topology. For all sequences identified via PSI-BLAST, we kept sequences with >6 (rather than 7) TM segments, as DeepTMHMM had previously failed to predict TM7 despite the presence of TM7 helices in the associated structural models ^10^. Finally, to reduce redundancy, and thus simplify computation and presentation, CD-HIT (https://cd-hit.org) ^55, 56^ was used to cluster sequences— first by 70% for the initial BLASTP sequence similarity network (**Figure 1H**), then by 50% for all subsequent analyses—keeping the longest sequence as the cluster representative.

A notable limitation of this approach is the use of metagenomics for the identification of some prokaryotic 7TMICs. As these data are assembled from environmental samples, these sequences could be misidentified. While this possibility cannot be completely discounted, it is not a compelling problem, as most of the metagenomically identified sequences described herein correspond to tens-to-hundreds of homologous proteins in closed prokaryotic genomes. The only obvious exceptions are the small number of archaeal and bacterial sequences most closely resembling Class-B Euk7TMICs.

### *Ab initio* protein folding and structural analyses

All monomer models were downloaded from the AlphaFold Protein Structure Database. Protein multimers (5 models each) were generated for *A. bakeri* Orco, the example 7TMICs in **Figure 2**, *Symbiodium natans* A0A812K102 (the most central node in the structural network, **Figure 3A**), and 7 additional structures derived from Foldseek clustering of 7TMICs by 50% alignment coverage (thus representing nearly the entire 7TMIC fold space, **Figure S2D** and supplemental data). Predictions were performed in Google Colaboratory (https://research.google.com/colaboratory) using AlphaFold2+MMSeqs2 as implemented by Colabfold (https://github.com/sokrypton/ColabFold) ^40–43^. These models were not interpreted as accurate predictions of protein stoichiometry, but rather as hypothetical tetramers and as indirect predictions of protein-protein interactions. We also generated hypothetical dimers, trimers, and pentamers for *A. bakeri* Orco (available in the supplemental data) and observed that the protein subunits assembled in a globally similar way – i.e. closest contact at the anchor domain(s). Transmembrane prediction was performed using DeepTMHMM, Phobius, and PeSTo webservers, as described above. Protein-protein interactions were predicted using a local implementation of PeSTo (https://github.com/LBM-EPFL/PeSTo). All proteins were visualized in PyMol. Visualized structural alignments were generated using C*oot* ^57^.

### Network and conservation analyses

We used graph-based strategies for visualizing relatedness among proteins ^58^. Structure-based networks were generated from the results of all-to-all DaliLite or Foldseek searches, where connections are derived from Z-scores >8 or TM scores >0.5, respectively. BLASTP sequence-based networking was performed using the CLANS webserver (https://toolkit.tuebingen.mpg.de/tools/clans) and a local implementation of CLANS ^52–54^, using attraction values derived from E-values <0.01; clusters were identified using the built-in network clustering algorithm with the global averages option.

PSI-BLAST networking was performed via all-to-all PSI-BLAST searches using a local implementation of BLAST+ ^36^. First, BLAST databases were prepared from the sequences databases described above. Insect Ors were excluded, as they are an insect-specific radiation ^28, 59^; their removal thus reduced the likelihood of spurious connectivity between distantly related 7TMICs, as demonstrated by their relatively high connectivity in the BLASTP network (see **Figure 1H**). In other words, the removal of Ors hypothetically weakened network connectivity overall, but increased our confidence in homology between linked sequences. BLAST+ was then used to perform all-to-all PSI-BLAST searches, stopping at either convergence or 10 iterations. PSSMs were generated with an E-value cutoff of 0.01 and the final network was assembled from hits where the PSSM query coverage was >70%. For any query-to-subject relationship, only the first significant PSI-BLAST hit was kept, corresponding to the weakest significant connection (as connections tend to strengthen in subsequent PSI-BLAST iterations), thus providing the most conservative interpretation of the network. The opsin control/outgroup databases were from previous studies ^60, 61^.

All networks were visualized, annotated, and quantitatively analyzed in CLANS, CytoScape ^62^ and Adobe Illustrator.

For conservation analyses, query-centered sequence alignments were first produced by Foldseek; in the figures, we visualized the alignment from the model with the highest closeness centrality (i.e. the centermost model; A0A812K102) from the structural similarity network. Amino acid sequence identity scores were calculated in Jalview. Embedding-based conservation scores were calculated using the esm2_t33_650M_UR50D protein language model ^63^, via the methods and scripts described by ^38^ (https://github.com/esbgkannan/kibby). The mean conservation scores were calculated by spline interpolating each individual data series (corresponding to each protein) to match the length of A0A812K102, then averaging those values; as such, family– and subfamily-specific conservation patterns are likely not represented in the average curve. The embedding-based conservation scores, PeSTo predictions, and query-centered multiple sequence alignments for all representative models are available in the supplemental data. Pearson’s correlation analysis (and the Bayesian equivalent) was performed in JASP.

### Phylogenetics

7TMIC GenBank accession numbers from our 50% clustered sequence database were matched to UniProt and 1947 AlphaFold-derived protein models were downloaded from the AlphaFold Protein Structure Database. All subsequent phylogenetic analyses were carried out on these 1947 representative proteins.

Muscle5 was used to generate the ensemble of multiple sequence alignments (MSAs) ^64^. Because the alignments were extremely long and gap rich (**Table S1**), MSAs were trimmed using trimal with the-gappyout option ^65^. Each trimmed MSA was then used to generate phylogenetic trees using FastTree2 ^66^, using 3 different amino acid substitution models (JTT, WAG, and LG), and with branch lengths rescaled to optimize the Gamma20 likelihood. The initial MSAs had extremely high dispersion and extreme lack of consensus (**Figure S3B**) indicating widespread alignment errors ^64^. These errors resulted in non-trivial topological differences in the phylogenetic trees, resulting in extreme non-consensus (even for obviously monophyletic clades, such as the insect Ors). This suggested a high degree of phylogenetic instability, likely due to both alignment errors (from low PID) and phylogenetic errors (e.g. long branch attraction).

To minimize alignment and phylogenetic errors, we repeated MSA and tree inference after identifying and removing rogue taxa (i.e. the most unstable leaves in the previous ensemble analysis) via RogueNaRok ^67^. Although the resulting ensemble of MSAs still had high dispersion, the resulting phylogenetic trees were more consistent in the assignment of the various subfamilies as monophyletic clades (**Figure S3C**). These trees were used for subsequent analysis.

The structural phylogeny was generated using fold_tree ^45^. Here, we emphasize the tree derived from all-against-all TM-scores, thereby sampling structural space based on pairwise global rigid structural comparisons, mirroring our network-based analysis, as described above. Structural trees derived from pairwise distances based on the Foldseek structural alphabet (**Figure S3F**) or pairwise lDDT scores (**Figure S3G**) produced radically different topologies; neither has obviously high congruence with the sequence-based phylogenetics, nor with the presumed taxonomy of the species included in this analysis. All trees were analyzed using the ape ^68^, phytools ^69^, and treespace ^70^ R packages. Tree topology space was explored by principal coordinate analysis of the Robison-Folds distances between the unrooted phylogenies. Trees were visualized and annotated using R, iTol (https://itol.embl.de/) ^71^, and Adobe Illustrator.

### Protein nomenclature

Most previous naming conventions have not been evolutionarily informed. Terms such as Gustatory receptor-like (Grl and GRL) do not refer to monophyletic clades, but instead correspond to many taxon-specific 7TMIC branches. For animal Grls and unicellular eukaryotic GRLs, the terms were chosen because they resembled insect Grs in either amino acid sequence and/or tertiary structure ^8,9,19^; by contrast, insect Grls were named based on the *absence* of sequence similarity to Grs despite the presence of structural similarity ^10^. While these terms are useful in situational contexts, they are uninformative at long evolutionary scales. We propose that the 7TMIC superfamily be split into domain-specific families; for eukaryotes, these are Class-A and Class-B. We suggest that the more complex nomenclature of previous work (e.g. Or, Gr, GrlHz) should be reserved for taxon-specific contexts. Relatedly, the evolution of Arch7TMICs and Bac7TMICs is highly reticulated. Although we saw proximity between Heimdallarchaeota 7TMICs and Euk7TMICs in our structure-based phylogeny, there were no other clear recapitulations of Asgard/Eukaryota monophyly or the Archaea-Bacteria split. Therefore, “Arch7TMIC” and “Bac7TMIC” serve only as terms of convenience, and we strongly caution that they do not refer to monophyletic clades.

## Supplemental Figures

**Figure S1.**
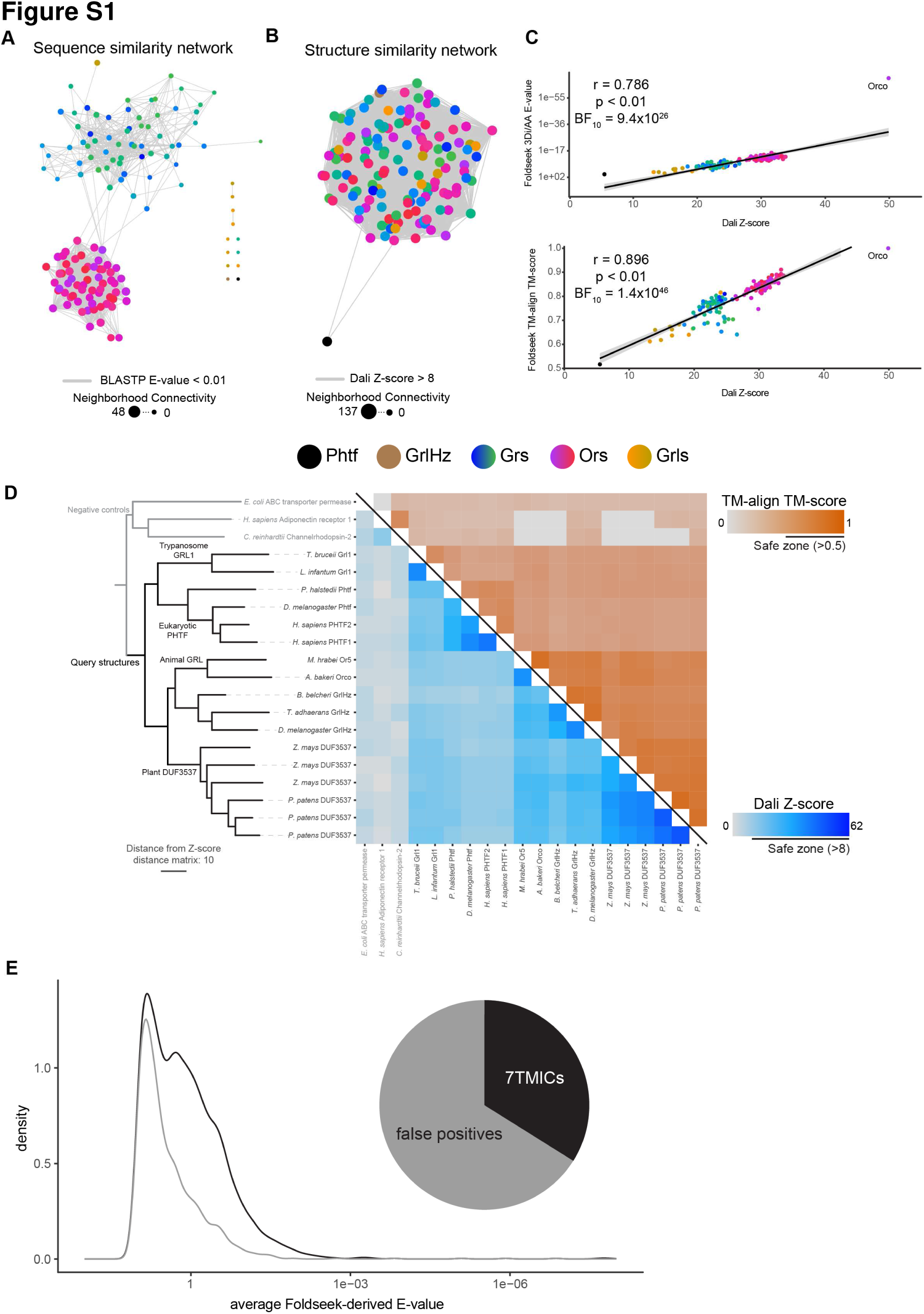
All-to-all pairwise protein similarity networks of *D. melanogaster* 7TMICs, Foldseek benchmarking, and summary of the Foldseek screen. (**A**) All-to-all BLASTP network of *D. melanogaster* 7TMICs; consistent with their low pairwise sequence similarity, this analysis fails to link every 7TMIC to all others. Rather, the major *D. melanogaster* classes (Ors and Grs) are separated into two identifiable community structures, with sparse connectivity among the Grs, and between the Grs and Ors. Other 7TMICs—including Grls, GrlHz, Phtf and two Grs—form singlets, indicating an inability to identify hypothetical homologs using BLASTP. (**B**) All-to-all Dali network of *D. melanogaster* 7TMICs. In contrast to (A), structural comparisons result in a “hairball” structure, wherein nearly all proteins are linked to all others, excepting Phtf, which is presumed to be the most distantly related. (**C**) Plots of structural similarity scores between Orco and other *D. melanogaster* 7TMICs, comparing Dali to Foldseek-derived scores. Foldseek generates Orco-to-all E-values that tightly correlate with the rapidly generated 3Di+AA-derived E-values (top) and the slowly generated TM-align-derived TM-scores (bottom). (**D**) Protein models used in the Foldseek screen, and negative controls used for subsequent Dali-based validation, with a clustering dendrogram based on all-to-all Dali comparisons between the queries and negative controls. The dendrogram is derived from the Dali Z-score distance matrix. The heatmap shows all-to-all Dali Z-scores and TM-scores. (**E**) Stacked density plot showing the frequency distribution of the hits of the Foldseek screen, by E-value, with the inset pie-chart showing the proportion of true positives to false positives. Most true positives had relatively poor E-values, with similar or worse scores than many false positives, demonstrating the need for structural validation in a Foldseek screen.

**Figure S2.**
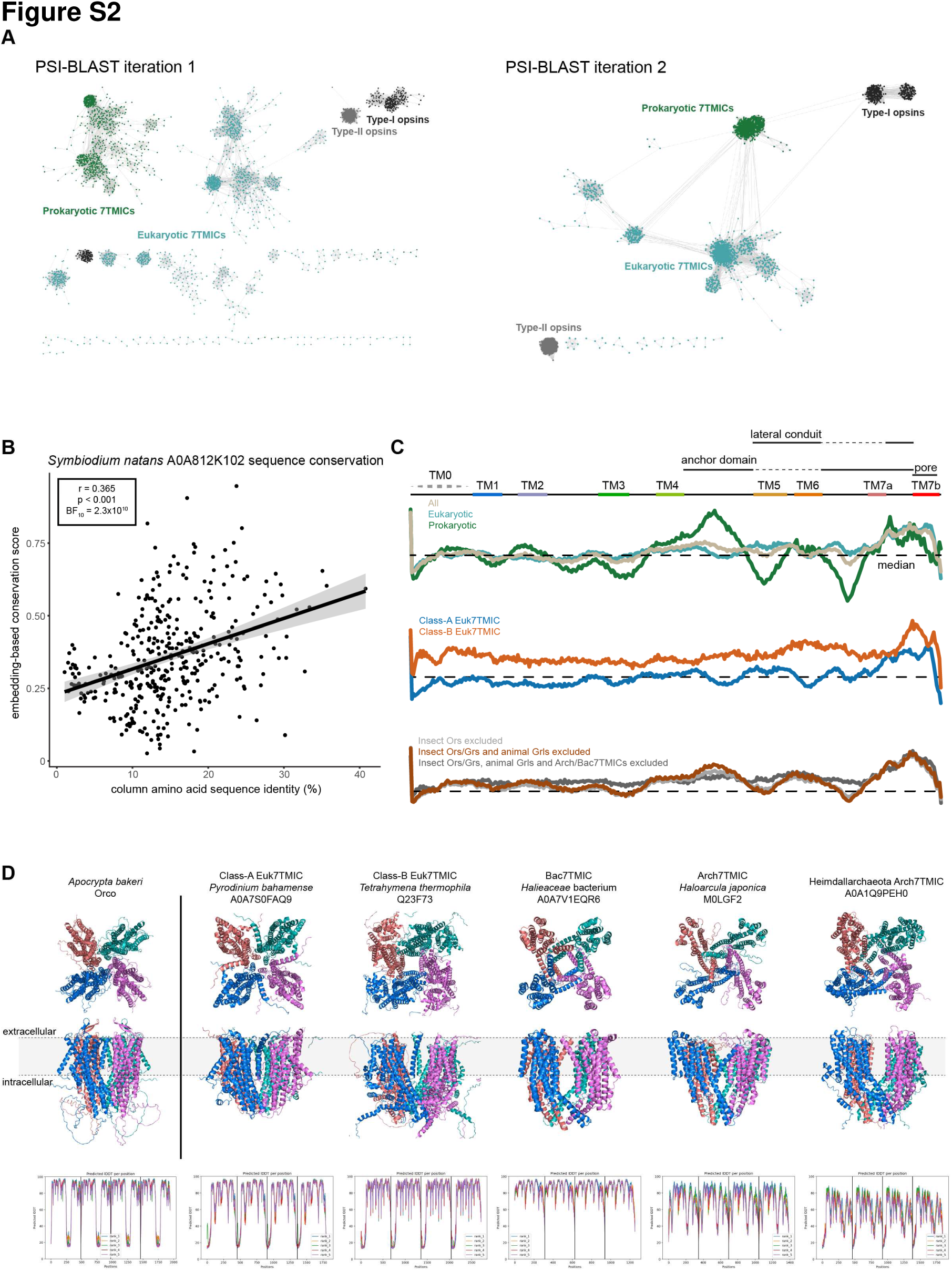
Initial iterations of the PSI-BLAST sequence similarity networks, 7TMIC sequence conservation analysis, and predicted quaternary structures of select, newly-identified 7TMICs. (**A**) Sequence similarity networks were generated by all-to-all PSI-BLAST searches of a 50% clustered sequence database of 7TMICs, alongside databases of Type-I and Type-II opsins. Iterations 1 and 2 are visualized here. Subsequent iterations resemble the clustering pattern of iteration 3, as visualized in Figure 3, albeit with strengthening community structures. Left: PSI-BLAST iteration 1. In this network, sequences formed several non-contiguous clusters, and failed to cluster together 7TMICs and Type-I opsins, which is expected given the substantial sequence dissimilarity of 7TMICs. Right: PSI-BLAST iteration 2. Surprisingly, PSI-BLAST networking produced bidirectional linking of the majority of 7TMICs, although presumed spurious linkages to outgroups began to form (which did not greatly multiply in subsequent iterations), and a small number of 7TMICs do not form links to the core 7TMIC cluster(s) (although all join a 7TMIC community structure by iteration 3 (Figure 3B)). (**B**) Embedding-based conservation scores weakly but significantly correlate with column sequence identity from the A0A812K102-centered sequence alignment. (**C**) Average embedding-based conservation scores for different subsets of 7TMICs, demonstrating that, while family-specific patterns exist, the conservation of anchor domain and pore regions is consistent. The TM and domain labels are derived from A0A812K102, as visualized in Figure 3. (**D**) Predicted tetramers for select 7TMICs. Top: top (presumed extracellular) and side views of the tetrameric arrangement of 7TMICs predicted by AlphaFold-Multimer, showing the formation of a hypothetical pore along TM7b, similar to *A. bakeri* Orco (far-left). Bottom: local Distance Difference Test (lDDT) scores (used to assess model confidence), plotted for each of the 5 replicate models generated. Each color represents a different replicate. Vertical black lines separate each of the modelled subunits. Generally, the transmembrane-spanning alpha helices are the most confidently predicted, leading to the similar patten of lDDT peaks and troughs across models.

**Figure S3.**
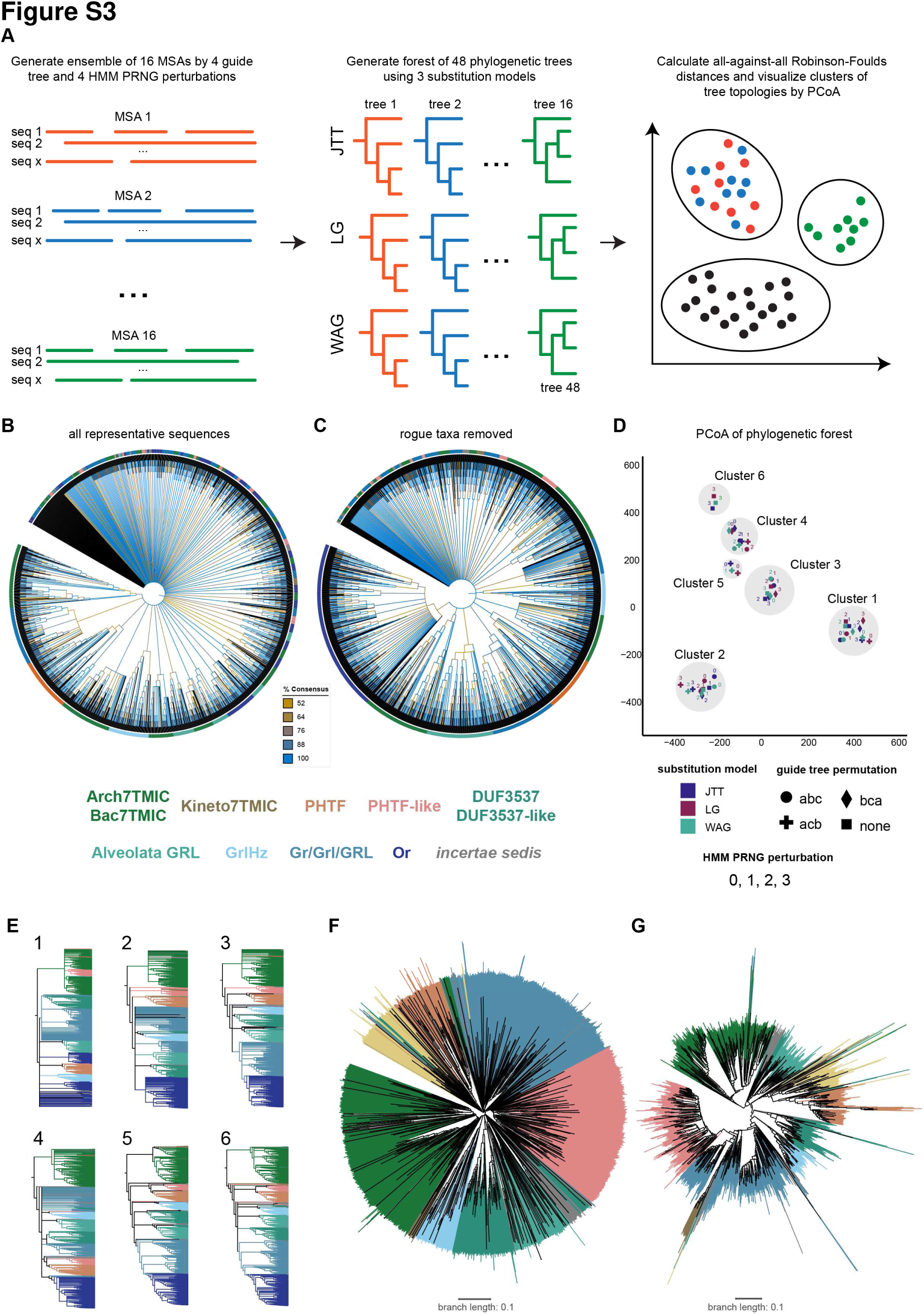
Phylogenetic and tree space analysis. (**A**) Pipeline for sequence-based phylogenetic analysis. First, an ensemble of 16 multiple sequence alignments (MSAs) are made by perturbating the guide tree and the Hidden Markov model’s pseudorandom number generator (HMM PRNG). Second, phylogenetic trees are generated for each of the MSAs, using 3 different amino acid substitution models, resulting in 48 trees. Finally, differences in the topology of the 48 trees are calculated by pairwise Robinson-Foulds distances; the resulting distance matrix is subsequently visualized in two dimensions by principal coordinate analysis (PCoA). (**B**) Majority consensus tree for the 48 phylogenetic trees based on alignments of the representative 7TMIC sequences. 7TMIC clades/colors were assigned manually based on visual inspection of a CLANS-based clustering analysis. Branch colors indicate the percent consensus. There is essentially no clear consensus among these 48 initial trees; obviously monophyletic clades—such as insect Ors— are not reliably predicted, suggesting substantial alignment/phylogenetic errors (as expected for this highly divergent superfamily). (**C**) Majority consensus tree for the 48 phylogenetic trees based on alignments of a 7TMIC dataset where rogue taxa (i.e. the most phylogenetically unstable leaves) have been removed (with colors matching (B)). While there is still no greatly informative majority consensus topology, this analysis better recapitulates more obvious monophyletic clades, with higher branch consensus, indicating that errors have been minimized (but not eliminated, which we did not expect to occur at these levels of sequence dissimilarity). (**D**) PCoA of Robinson-Foulds tree space for trees from (C). Trees form 6 topology clusters. (**E**) Majority consensus trees for each of the 6 clusters, with colors matching (C) and (D). Five of these clusters agree that Kineto7TMICs branch proximally to prokaryotic 7TMICs, consistent with the hypothesis that kinetoplastids (and allies: Discoba) split early in eukaryotic evolution ^44^. Clusters 1 and 4 do not have majority consensus on deep 7TMIC branching. The remaining clusters suggest there are at least two Euk7TMIC families, termed Class-A and Class-B Euk7TMICs, but do not agree on the monophyly of Class-B Euk7TMICs. Clusters 4-6 suggest Class-B monophyly, while clusters 1-3 suggest that many proteins are basally branching (and thus, paraphyly). Given that structure-based phylogenetics suggest a monophyletic Class-B, this discordance may be the result of lingering long branch attraction or other errors resulting from the inclusion of rapidly evolved, horizontally-transferred, or structurally-convergent proteins. (**F**) Structural phylogeny derived from pairwise distances used the Foldseek 3Di structural alphabet, with colors matching the panels above. This tree is presented as rooted, but as in Figure 4, the true root is likely within the prokaryotic 7TMICs, at the location of the Last Universal Common Ancestor. (**G**) Structural phylogeny derived from pairwise lDDT scores, with colors matching the panels above. As in (F), the true root is likely at location of the Last Universal Common Ancestor.

## Supplemental Table

**Table S1.** Muscle5 multiple sequence alignment analysis. Column confidence is a measure of the reproducibility of each column, where 0 indicates the column is never found, and 1 indicates it is found across all alignments. Dispersion is measured as the median dispersion of aligned letter pairs over the ensemble (D_LP), and the median dispersion of columns over the ensemble (D_Cols) (Robert Edgar, personal communication, 10 May 2023), where 0 is all the same and 1 is all different. Dispersion was extremely high. For the initial set of alignments: D_LP=0.5836 D_Cols=1.0000. After removal of rogue taxa: D_LP=0.5855 D_Cols=1.0000.

| MSA Replicate | MSA Perturbations |  | All Sequences |  | No Rogue Taxa |  |
| --- | --- | --- | --- | --- | --- | --- |
|  | Guide Tree | PRNG | Columns | Column Confidence | Columns | Column Confidence |
| 1 | abc | 0 | 38154 | 0.315 | 36865 | 0.311 |
| 2 | abc | 1 | 38357 | 0.297 | 34833 | 0.295 |
| 3 | abc | 2 | 36970 | 0.314 | 35376 | 0.325 |
| 4 | abc | 3 | 37064 | 0.314 | 31247 | 0.304 |
| 5 | acb | 0 | 38375 | 0.31 | 35627 | 0.304 |
| 6 | acb | 1 | 40342 | 0.298 | 34754 | 0.302 |
| 7 | acb | 2 | 37080 | 0.317 | 34994 | 0.321 |
| 8 | acb | 3 | 35735 | 0.313 | 31345 | 0.305 |
| 9 | bca | 0 | 38391 | 0.315 | 35914 | 0.314 |
| 10 | bca | 1 | 39831 | 0.3 | 35813 | 0.294 |
| 11 | bca | 2 | 37647 | 0.316 | 34847 | 0.322 |
| 12 | bca | 3 | 36515 | 0.305 | 31406 | 0.303 |
| 13 | none | 0 | 38700 | 0.3 | 35842 | 0.309 |
| 14 | none | 1 | 40114 | 0.306 | 34808 | 0.293 |
| 15 | none | 2 | 37443 | 0.308 | 34932 | 0.309 |
| 16 | none | 3 | 37831 | 0.309 | 32689 | 0.298 |

