## Supplementary material for "Remote homolog detection places insect chemoreceptors in a cryptic protein superfamily spanning the tree of life": Key resource table

| **REAGENT or RESOURCE** | **SOURCE** | **IDENTIFIER** |
| --- | --- | --- |
| **Deposited data** | | |
| All data | this paper | <https://doi.org/10.5061/dryad.fqz612jz9> |
| **Software and algorithms** | | |
| AlphaFold Protein Sequence Database | (Jumper et al., 2021; Varadi et al., 2022) | [https://alphafold.ebi.ac.uk](https://alphafold.ebi.ac.uk/) |
| ape | (Paradis & Schliep, 2019) | http://ape-package.ird.fr |
| BLAST+ | (Altschul et al., 1997) | [https://www.ncbi.nlm.nih.gov](https://www.ncbi.nlm.nih.gov/) |
| CD-HIT | (Fu et al., 2012; Li & Godzik, 2006) | [https://cd-hit.org](https://cd-hit.org/) |
| CLANS | (Frickey & Lupas, 2004; Gabler et al., 2020; Zimmermann et al., 2018) | <https://toolkit.tuebingen.mpg.de/tools/clans> |
| ColabFold | (Mirdita et al., 2017, 2019, 2022; Mitchell et al., 2019) | <https://github.com/sokrypton/ColabFold> |
| C*oot* | (Emsley et al., 2010) | [https://www2.mrc-lmb.cam.ac.uk/personal/pemsley/coot](https://www2.mrc-lmb.cam.ac.uk/personal/pemsley/coot/) |
| Cytoscape | (Shannon et al., 2003) | [https://cytoscape.org](https://cytoscape.org/) |
| DeepTMHMM | (Hallgren et al., 2022) | [https://dtu.biolib.com/DeepTMHMM](https://dtu.biolib.com/DeepTMHMM/) |
| esm2_t33_650M_UR50D | (Lin et al., 2023) | [https://huggingface.co/facebook/esm2_t33_650M_UR50D](https://huggingface.co/facebook/esm2_t33_650M_UR50D/) |
| FastTree 2.1 | (Price et al., 2010) | [http://www.microbesonline.org/fasttree](http://www.microbesonline.org/fasttree/) |
| fold_tree | (Moi et al., 2023) | <https://github.com/DessimozLab/fold_tree> |
| Foldseek | (van Kempen et al., 2023) | https://github.com/steineggerlab/foldseek |
| Illustrator | Adobe Inc. | <https://www.adobe.com/products/illustrator.html> |
| iTol | (Letunic & Bork, 2021) | [https://itol.embl.de](https://itol.embl.de/) |
| JASP | JASP Team | [https://jasp-stats.org](https://jasp-stats.org/) |
| Muscle5 | (Edgar, 2022) | [https://www.drive5.com/muscle](https://www.drive5.com/muscle/) |
| PeSTo | (Krapp et al., 2023) | [https://github.com/LBM-EPFL/PeSTo](https://github.com/LBM-EPFL/PeSTo/) |
| Phobius | (Käll et al., 2004, 2007) | [https://phobius.sbc.su.se](https://phobius.sbc.su.se/) |
| phytools | (Revell, 2012) | [https://github.com/liamrevell/phytools](https://github.com/liamrevell/phytools/) |
| PyMol | Schrödinger, LLC. | [https://pymol.org](https://pymol.org/) |
| RogueNaRok | (Aberer et al., 2013) | [https://github.com/aberer/RogueNaRok](https://github.com/aberer/RogueNaRok/) |
| R studio | R Studio Team | [http://www.rstudio.com](http://www.rstudio.com/) |
| Sequence conservation scripts | (Yeung et al., 2023) | <https://github.com/esbgkannan/kibby> |
| treespace | (Jombart et al., 2017) | <https://github.com/thibautjombart/treespace> |
| trimal | (Capella-Gutiérrez et al., 2009) | <http://trimal.cgenomics.org/trimal> |
